# The molecular principles underlying diverse functions of the SLC26 family of proteins

**DOI:** 10.1101/2023.12.10.570988

**Authors:** Satoe Takahashi, Kazuaki Homma

## Abstract

Mammalian SLC26 proteins are membrane-based anion transporters that belong to the large SLC26/SulP family, and many of their variants are associated with hereditary diseases. Recent structural studies revealed a strikingly similar homodimeric molecular architecture for several SLC26 members, implying a shared molecular principle. Now a new question emerges as to how these structurally similar proteins execute diverse physiological functions. In this study we sought to identify the common vs. distinct molecular mechanism among the SLC26 proteins using both naturally occurring and artificial missense changes introduced to SLC26A4, SLC26A5, and SLC26A9. We found: (i) the basic residue at the anion binding site is essential for both anion antiport of SLC26A4 and motor functions of SLC26A5, and its conversion to a nonpolar residue is crucial but not sufficient for the fast uncoupled anion transport in SLC26A9; (ii) the conserved polar residues in the N- and C-terminal cytosolic domains are likely involved in dynamic hydrogen-bonding networks and are essential for anion antiport of SLC26A4 but not for motor (SLC26A5) and uncoupled anion transport (SLC26A9) functions; (iii) the hydrophobic interaction between each protomer’s last transmembrane helices, TM14, is not of functional significance in SLC26A9 but crucial for the functions of SLC26A4 and SLC26A5, likely contributing to optimally orient the axis of the relative movements of the core domain with respect to the gate domains within the cell membrane. These findings advance our understanding of the molecular mechanisms underlying the diverse physiological roles of the SLC26 family of proteins.

## INTRODUCTION

The solute carrier 26 (SLC26)/sulfate permease (SulP) proteins constitute a large gene family, many of which function as solute transporters (Alper and Sharma, 2013). Increasing number of genetic variants are associated with various human diseases such as nephrocalcinosis (SLC26A1), hyperoxalemia (SLC26A1), diastrophic dysplasia (SLC26A2), achondrogenesis (SLC26A2), atelosteogenesis (SLC26A2), multiple epiphyseal dysplasia (SLC26A2), congenital chloride diarrhea (SLC26A3), deafness (SLC26A4 and SLC26A5), bicarbonate metabolism-related diseases (SLC26A6), hypothyroidism (SLC26A7), asthenozoospermia (SLC26A8), bronchiectasis (SLC26A9), and dysregulation of chloride homeostasis and neuroactivity (SLC26A11) (Alper and Sharma, 2013; Li et al., 2023; Stenson et al., 2020). The amino acid sequences among the SLC26 family members are quite similar, and recent structural studies revealed that their overall molecular architectures are also similar for mammalian SLC26A4 (Liu, 2023), SLC26A5 (Bavi et al., 2021; Butan et al., 2022; Futamata et al., 2022; Ge et al., 2021), SLC26A6 (Tippett et al., 2023), and SLC26A9 (Chi et al., 2020; Walter et al., 2019). Thus, it is conceivable that a common molecular mechanism underlies their diverse physiological roles, and the pathogenic variants found in one SLC26 protein may affect the functions of the other family members in similar manners.

However, it is not clear how SLC26 family members with similar structures can support their diverse physiological functions. It is possible that certain molecular features are not shared among family members and thus some variants are pathogenic only in a certain family member(s) but not in others. In this study, we sought to understand the common vs. distinct molecular mechanisms of the SLC26 proteins. To this end, we focus on three mammalian SLC26 family members with highly distinct functionalities, SLC26A4, SLC26A5, and SLC26A9. SLC26A4 (pendrin) is an electroneutral coupled anion exchanger that is essential for normal inner ear function (Choi et al., 2011; Everett et al., 2001; Everett et al., 1997; Kim and Wangemann, 2010; Li et al., 2013). SLC26A5 (prestin) is a voltage-driven motor protein responsible for cochlear amplification and thus essential for normal hearing (Cheatham et al., 2004; Dallos et al., 2008; Liberman et al., 2002; Wu et al., 2004; Zheng et al., 2000). SLC26A5 also reportedly retains small anion exchanging function (Mistrik et al., 2012) but with minimal physiological significance (Takahashi et al., 2023). SLC26A9 mediates coupled or uncoupled anion transport (Chang et al., 2009b; Dorwart et al., 2007; Loriol et al., 2008; Walter et al., 2019; Xu et al., 2005). The uncoupled chloride transport is rapid and shows discrete channel-like unitary conductance (Chang et al., 2009b). SLC26A9 plays crucial roles in gastric acid secretion (Xu et al., 2008) and airway clearance (Anagnostopoulou et al., 2012). The roles of SLC26A9 in renal chloride excretion and arterial pressure regulation are also reported (Amlal et al., 2013) despite its low expression in the kidney (Chang et al., 2009b; Xu et al., 2005).

We examined the functional consequences of various missense changes introduced to these three SLC26 proteins. We found that (i) the basic residues at the anion substrate binding site are crucial for the anion antiport and voltage-dependent motor functions of SLC26A4 and SLC26A5, respectively, whereas nonpolar residue at the equivalent site is crucial but not sufficient for rapid uncoupled electrogenic anion transport in SLC26A9, that (ii) conserved polar residues at the interface between the N- and C-terminal cytosolic domains likely contribute to form dynamic hydrogen-bonding networks and are crucial for the functions of SLC26A4, but not for SLC26A5 and SLC26A9, and that (iii) the hydrophobic interaction at the dimerization interface at the C-terminal ends of the last transmembrane helix, TM14, is not of functional importance for uncoupled anion transport in SLC26A9 but essential for the anion antiport and motor functions of SLC26A4 and SLC26A5, respectively. This comparative functional study thus provides mechanistic insights underlying the diverse physiological functions of the SLC26 family of proteins, which refines the accuracy of our pathogenicity assessment of disease-associated variants in this important family of proteins.

## RESULTS

### SLC26 constructs generated in this study

Recent single particle cryo-EM studies have revealed homodimeric structures for mammalian SLC26A4 (Liu, 2023) (**Fig. 1**, left), SLC26A5 (Bavi et al., 2021; Butan et al., 2022; Futamata et al., 2022; Ge et al., 2021) (**Fig. 1**, middle), SLC26A6 (Tippett et al., 2023), and SLC26A9 (Chi et al., 2020; Walter et al., 2019) (**Fig. 1**, right). The N- and C-termini physically interact to one another to form the cytoplasmic domain (CD), and the intervening sequence constitutes the transmembrane domain (TMD). TMD is comprised of fourteen transmembrane helices (TM1-14) that are divided into the core (TM1-4, TM8-11) and gate (TM5-7, TM12-14) subdomains. Each SLC26 protomer has its own anion binding pocket and anion translocation pathway between these two TMD subdomains. Basic residues are found at the anion binding site except for SLC26A9 (**Fig. 1B**). Elevator-like up and down movements of the core domain with respect to the gate domain underlie the anion transport functions and are also thought to account for the motor function of SLC26A5(Kuwabara et al., 2023). Homodimerization appears to be essential for rigidly maintaining the axis of the elevator-like core domain movements perpendicular to the cell membrane plane. Two hydrophobic residues at the C-terminus of TM14 are conserved (**Fig. 1B**) and seem to contribute to homodimerization. They also seem to be crucial for determining and maintaining the orientation of the gate domains within the cell membrane (**Figs. 1A and 1C**). The N- and C-termini are interwound, and their interaction is likely stabilized by the hydrogen bond (H-bond) networks (**Figs. 1A and 1C**). Although conformational changes at this N- and C-terminal interaction site are subtle (Bavi et al., 2021; Ge et al., 2021; Liu et al., 2023), it is conceivable that the H-bond networks are dynamic and intimately linked to large conformational changes in TMD (**Figs. 1D and S1**), possibly contributing to inter-protomer functional communications.

**Figure 1.**
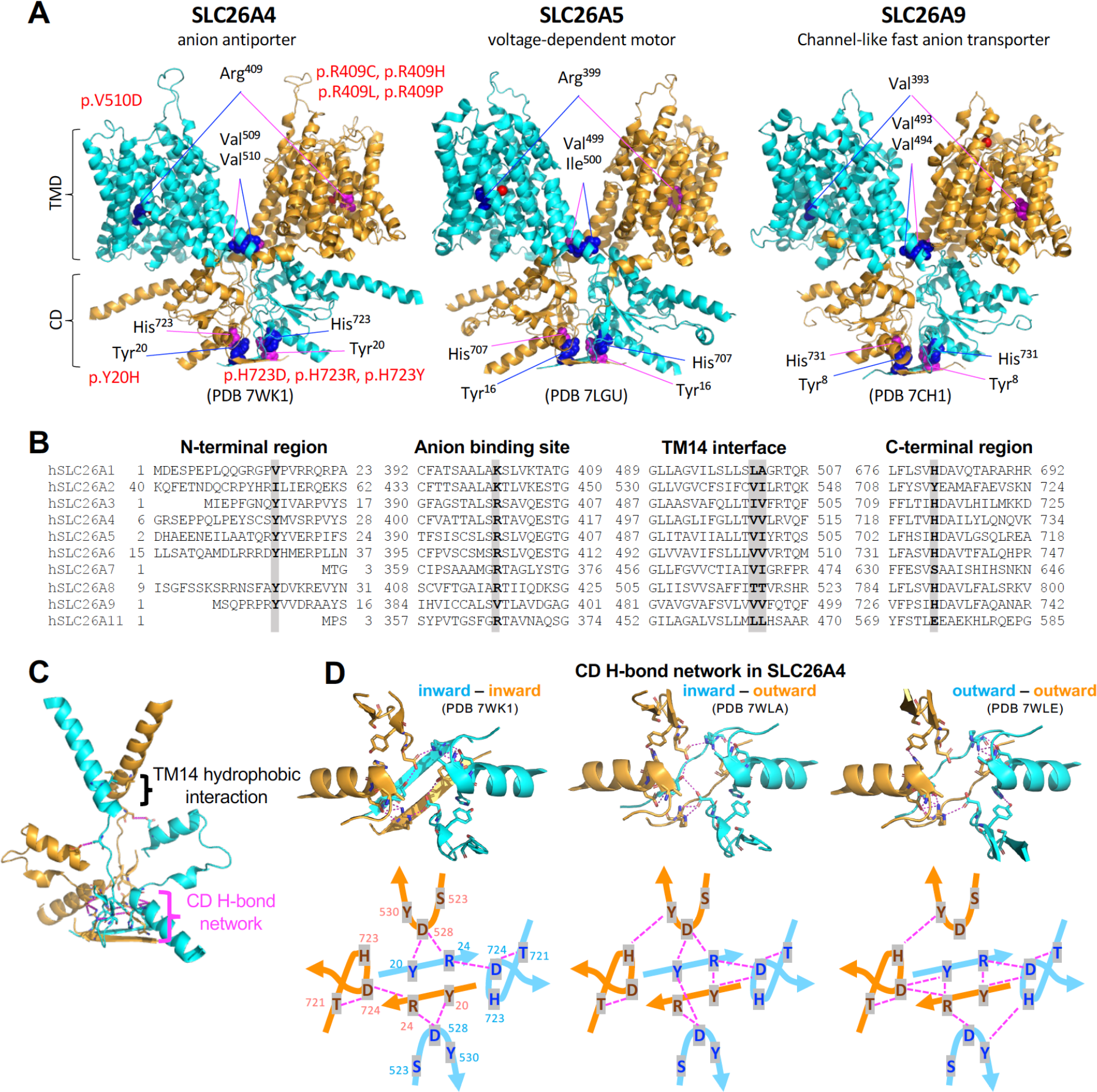
The amino acid residues of interest in this study. (**A**) The cryo-EM homodimeric structures of mouse SLC26A4 (left), human SLC26A5 (middle), and human SLC26A9 (right). The locations of the amino acid residues of interest in this study are shown in sphere representation in blue (in one protomer shown in cyan) and magenta (in the other protomer shown in orange). Missense variants shown in red were identified in *SLC26A4* in human patients. TMD, transmembrane domain; CD, cytoplasmic domain. (**B**) Amino acid sequence comparison among human SLC26 proteins for selected protein regions. The amino acid residues of interest in this study are indicated in bold and highlighted. The residue numbers at the N- and C-termini are also shown. (**C**) TM14 helices and selected C-terminal structures in mouse SLC26A4 (PDB: 7WK1). Tyr^20^, Arg^24^, Val^509^, Val^510^, Ser^517^, Ser^523^, Asp^528^, Tyr^530^, Asp^698^, Thr^721^, His^723^, and Asp^724^ are shown in stick representation. Magenta dashed lines (seen more easily in panel D) indicate potential hydrogen bond interactions. (**D**) The N- and C-terminal interaction in mouse SLC26A4 in three distinct structural modes as indicated. Tyr^20^, Arg^24^, Ser^523^, Asp^528^, Tyr^530^, Thr^721^, His^723^, and Asp^724^ are shown in stick representation. Magenta dashed lines indicate potential hydrogen bond interactions, which are schematically shown in lower panels for clarity.

The functional importance of these key regions of interest is implied by the presence of multiple disease-associated *SLC26A4* missense variants identified in humans (**Fig. 1A**, left). Hence, we systematically examined the functional consequences of the following missense changes, aiming at gaining mechanistic insights underlying the diverse functions of the SLC26 proteins: Y20H, V509G, V510D, V510I, H723D, and H723Y in SLC26A4; Y16H, R399V, I500V, and H707R in SLC26A5; Y8H, V393R, V493G, and H731R in SLC26A9. Among these, Y20H (c.58T>C), V510D (c.1529T>A), H723D (c.2167C>G), and H723Y (c.2167C>T) were identified in *SLC26A4* in human patients (Dai et al., 2008; Jang et al., 2014; Miyagawa et al., 2014; Zhao et al., 2014), while the others are not. All SLC26 constructs including WT controls were C-terminally tagged with mTurquoise2 (mTq2) and expressed in HEK293T cells in a doxycycline (Dox)-inducible manner for functional assays as in previous studies (Kuwabara et al., 2018; Wasano et al., 2020). To quantitatively assess membrane targeting, HEK293T cells heterologously expressing WT and mutated SLC26 proteins were treated with membrane impermeable sulfo-Cyanine3 (Cy3) NHS ester that labeled proteins expressed in the plasma membrane. The cells were subsequently lysed in a solution containing a mild detergent, and the solubilized SLC26 proteins were immunoprecipitated using nanobody that recognizes mTq2. The ratios of Cy3 and mTq2 fluorescence on beads, F_Cy3_/F_mTq2_, indicate the membrane targeting efficiencies of the SLC26 constructs (Futamata et al., 2022). HEK cells expressing only the mTq2 moiety were used as negative control. The F_Cy3_/F_mTq2_ ratios were normalized to WT controls and summarized in **Fig. 2**. We found that the F_Cy3_/F_mTq2_ ratios of all SLC26 constructs were higher than those of the negative controls (mTq2 alone), affirming that none of the missense changes introduced abrogated the membrane targeting of the SLC26 proteins. The relative F_Cy3_/F_mTq2_ ratios (**Fig. 2**) were used for correcting the measured anion transport activities of SLC26A4 and SLC26A9 (see below).

**Figure 2.**
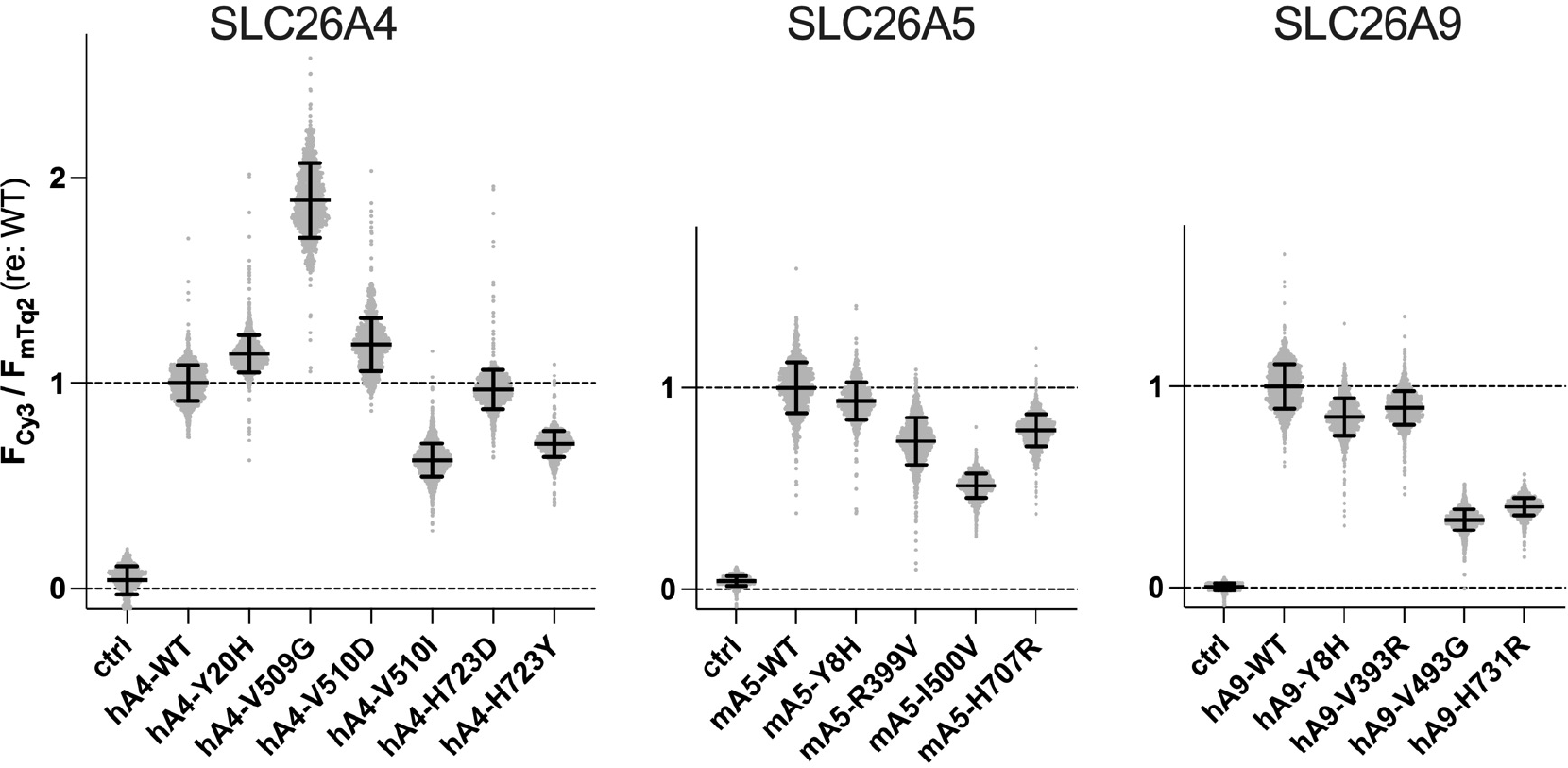
The membrane targeting efficiencies of the SLC26 constructs. HEK293T cells expressing various mTurquoise2 (mTq2)-tagged SLC26 proteins (hA4, human SLC26A4; mA5, mouse SLC26A5; and hA9, human SLC26A9) were treated with a membrane impermeable sulfo-Cy3 NHS ester to only label cell surface proteins. After lysing cells, mTq2-tagged SLC26 proteins were collected regardless of their original subcellular localizations using affinity beads that capture mTq2, and the ratios of Cy3 and mTq2 fluorescence, F_Cy3_ /F_mTq2_, were determined and compared to those of WTs. The results were normalized to the WTs. Error bars indicate propagated errors computed using the standard deviations of the F_Cy3_ /F_mTq2_ data. HEK cells expressing only the mTq2 moiety were used as negative control (ctrl).

### The anion binding site

Basic residues at the N-terminus of TM10 in the core domain (**Figs. 1A and 1B**) likely contribute to transiently attract and hold anion substrates during the transport process. The importance of this highly conserved basic residue in SLC26A4 (Arg^409^) is reflected by the presence of multiple disease-associated missense variants, i.e., R409C (c.1225C>T) (Chen et al., 2007), R409H (c.1226G>A) (Van Hauwe et al., 1998), R409L (c.1226G>T) (Chai et al., 2013), and R409P (c.1226G>C) (Park et al., 2003) and has been experimentally demonstrated (Dossena et al., 2009; Gillam et al., 2005; Wasano et al., 2020; Yuan et al., 2012; Zhang et al., 2022). In SLC26A6, Arg^404^ is experimentally demonstrated to be important for its anion antiport function (Tippett et al., 2023). Interestingly, SLC26A9, which mediates channel-like fast uncoupled electrogenic chloride transport (Chang et al., 2009b; Dorwart et al., 2007; Loriol et al., 2008), has a nonpolar residue, Val^393^, at the equivalent position (**Fig. 1B**). To examine the functional importance of Val^393^ in SLC26A9, we measured whole-cell currents in HEK293T cells expressing WT and V393R SLC26A9 (hA9-WT and hA9-V393R, respectively, in **Fig. 3**).

**Figure 3.**
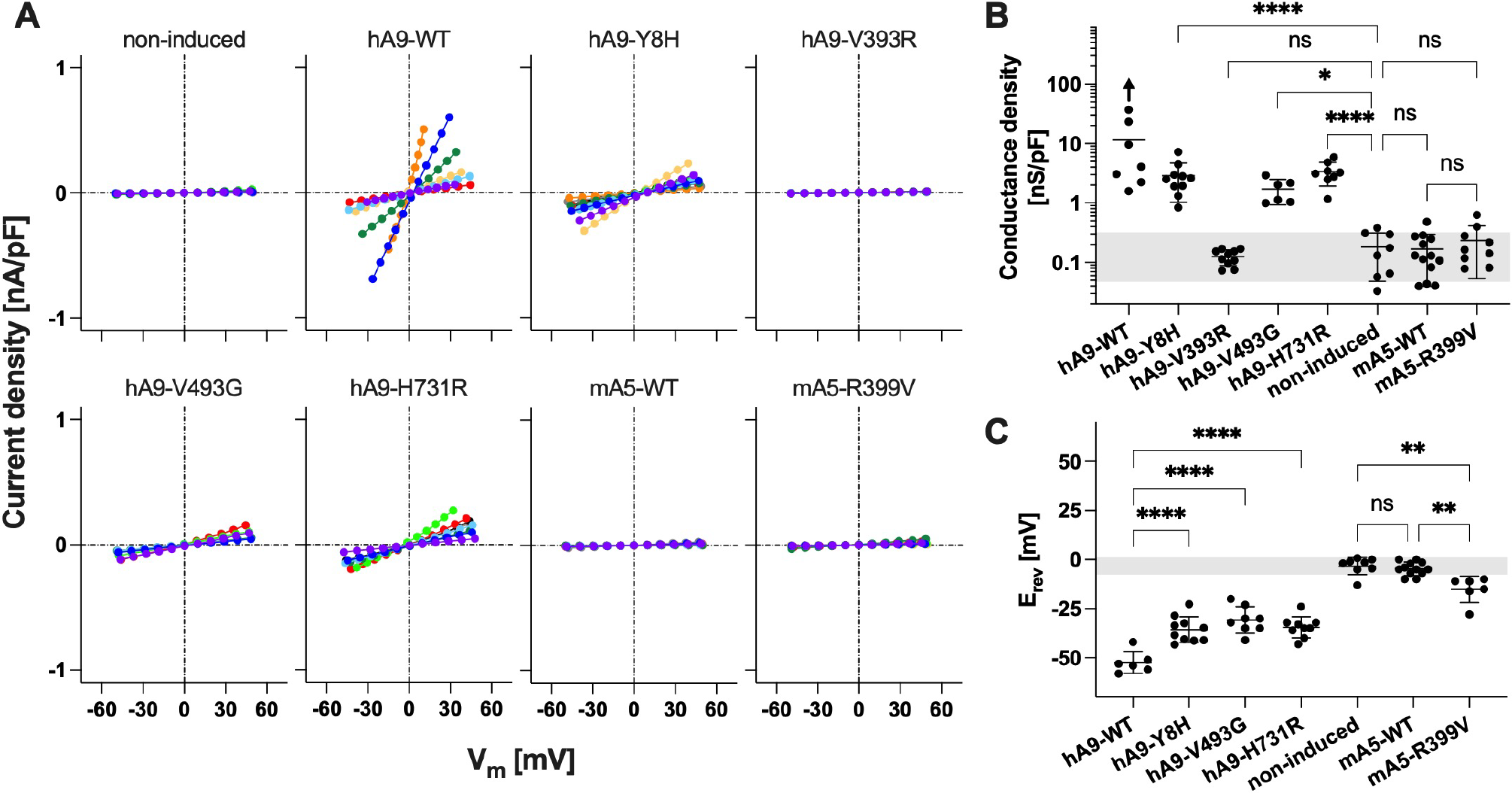
Whole-cell electrophysiological chloride transport assays. (**A**) Whole-cell conductance was recorded in HEK293T cells expressing various SLC26A9 and SLC26A5 proteins. The current-voltage data were corrected for the series resistance and divided by the cell membrane capacitance. Different colors indicate individual recordings. (**B**) Summaries of the whole-cell recordings. An upward arrow indicates underestimation of the mean conductance density for hA9-WT (see the main text). Error bars indicate standard deviations. A gray shade indicates mean ± standard deviation of non-induced negative control. (**C**) Summaries of reversal potentials (E_rev_) determined using a low chloride-containing intracellular solution. Error bars indicate standard deviation. A gray shade indicates mean ± standard deviation of non-induced negative control. In panels B and C, one-way ANOVA followed by Tukey’s post hoc test was performed. “ns”, *p* ≥ 0.05. “*”, 0.01≤ *p* < 0.05. “**”, 0.001≤ *p* < 0.01. “***”, 0.0001≤ *p* < 0.001. “****”, *p* < 0.0001.

Cells not treated with Dox (negative control) showed very small chloride current as expected, whereas those expressing WT (hA9-WT) showed very large currents (the whole-cell currents were corrected for cell size) (**Fig. 3A**). In some cells, the hA9-WT-mediated chloride conductance was so high that the measured whole-cell currents were limited by the series resistance of the recording pipette. Thus, the mean magnitude of the chloride conductance density shown in **Fig. 3B** for hA9-WT is underestimated to an unknown degree (indicated by an upward arrow). Conversion of Val^393^ to Arg (hA9-V393R) resulted in very small conductance density that was statistically indistinguishable from that of non-induced negative control (**Figs. 3A and 3B**). As expected, chloride conductance density was also small and indistinguishable from non-induced negative control in cells expressing SLC26A5 (mA5-WT, **Figs. 3A and 3B**). Interestingly, conversion of Arg^399^ to Val in SLC26A5 (mA5-R399V, **Fig. 3B**) slightly increased the mean conductance density although this increase was not statistically significant. We repeated whole-cell voltage clamp recordings using a low chloride-containing intracellular solution and determined the reversal potentials, E_rev_, that are V_m_ readings at zero current and are independent of the series resistance (thus are immune to uncertainty introduced by correction for the series resistance). As expected, cells expressing hA9-WT showed greatly hyperpolarized E_rev_ due to the large chloride conductance and the large inward transmembrane chloride gradient ([Cl^-^]_out_ = 148 mM, [Cl^-^]_in_ = 10 mM) (**Fig. 3C**). Slightly hyperpolarized E_rev_ with statistical significance was found in cells expressing mA5-R399V, suggesting that the elimination of the basic residue from the anion binding site conferred small uncoupled chloride transport function on SLC26A5.

This observation is in line with a recent report that R404V conferred a small, uncoupled chloride transport function on SLC26A6 (Tippett et al., 2023).

Since SLC26A9 also mediates iodide transport (Dorwart et al., 2007; Loriol et al., 2008), we further examined the transport function of WT and V399R SLC26A9 using iodide-sensitive mVenus^H148Q/I152L^ that was coexpressed with the SLC26A9 constructs (**Fig. 4A**). As described in detail in a preceding study (Wasano et al., 2020), this plate reader-based automated fluorometric assay also determined total expression of mTq2-tagged protein (F_mTq2_) and total cell density (OD_660nm_) (**Fig. 4B**). Combined with the membrane targeting efficiencies determined using sulfo-Cy3 NHS (**Fig. 2**, right panel), these extra measurements allowed determination of the amounts of mTq2-tagged SLC26 constructs expressed in the cell membrane, for which the measured iodide transport rates were corrected (**Fig. 4C**). As expected, cells expressing hA9-WT showed Dox-dependent increase in iodide transport activity, whereas those expressing hA9-V393R did not, further demonstrating the essentiality of Val^393^ for the anion transport function of SLC26A9.

**Figure 4.**
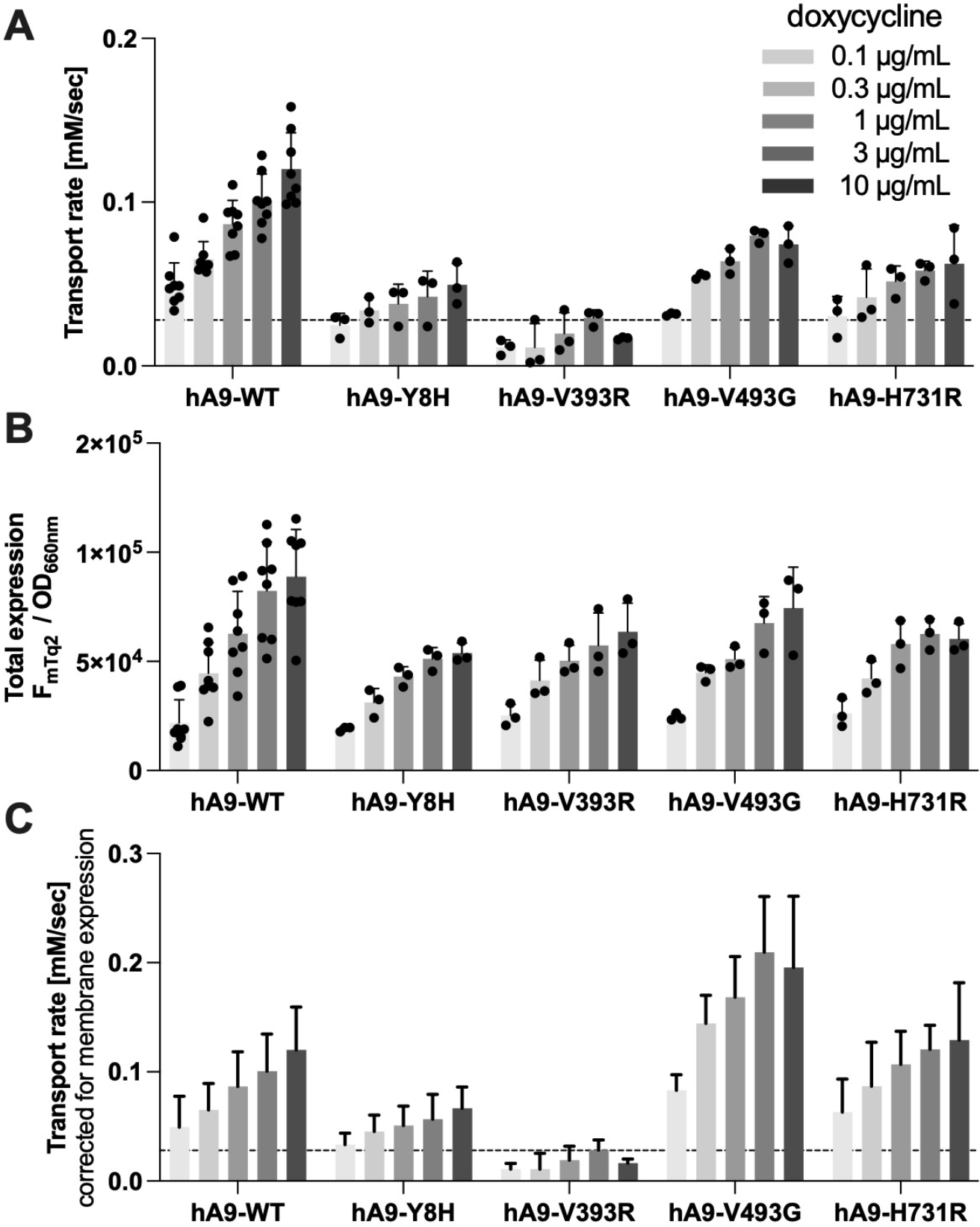
Multi-cell fluorometric iodide transport assays. (**A**) Iodide transport rates measured in HEK293T cells expressing various SLC26A9 constructs in a Dox-dependent manner. A horizontal dashed line indicates the basal iodide influx rate of non-induced cells (negative control). Error bars indicate standard deviation. (**B**) Ratios of mTq2 fluorescence (F_mTq2_) and the optical densities (OD_660nm_) of the cells expressing the mTq2-tagged SLC26A9 constructs, which quantify the total expressions of the SLC26A9 constructs. Error bars indicate standard deviation. (**C**) SLC26A9-mediated iodide transport rates corrected for the relative membrane expressions. Error bars indicate propagated errors computed from the standard deviations in panel A and the errors in the relative membrane expressions.

Although the anion transport by mammalian SLC26A5 is of minimal physiological significance (Takahashi et al., 2023), anions are still needed by SLC26A5 as extrinsic cofactors for supporting the fast motor function (Oliver et al., 2001; Santos-Sacchi et al., 2006). Arg^399^ very likely contributes to anion binding in SLC26A5, and its functional importance was experimentally demonstrated (Bai et al., 2009; Gorbunov et al., 2014). In line with these preceding studies, we found that HEK293T cells expressing R399V SLC26A5 (mA5-R399V) do not exhibit nonlinear capacitance (NLC), which is the electrical signature of the voltage-driven motor function (electromotility) of SLC26A5 (**Fig. 5A**, top right) despite only a small negative effect of R399V on membrane targeting (**Fig. 2**). We also found that conversion of Val^393^ to Arg does not confer NLC on SLC26A9 (hA9-V393R), at least in the experimentally testable voltage range (**Fig. 5A**, bottom right). Note that NLC measurement was not feasible in cells expressing hA9-WT due to excessively high membrane conductance.

**Figure 5.**
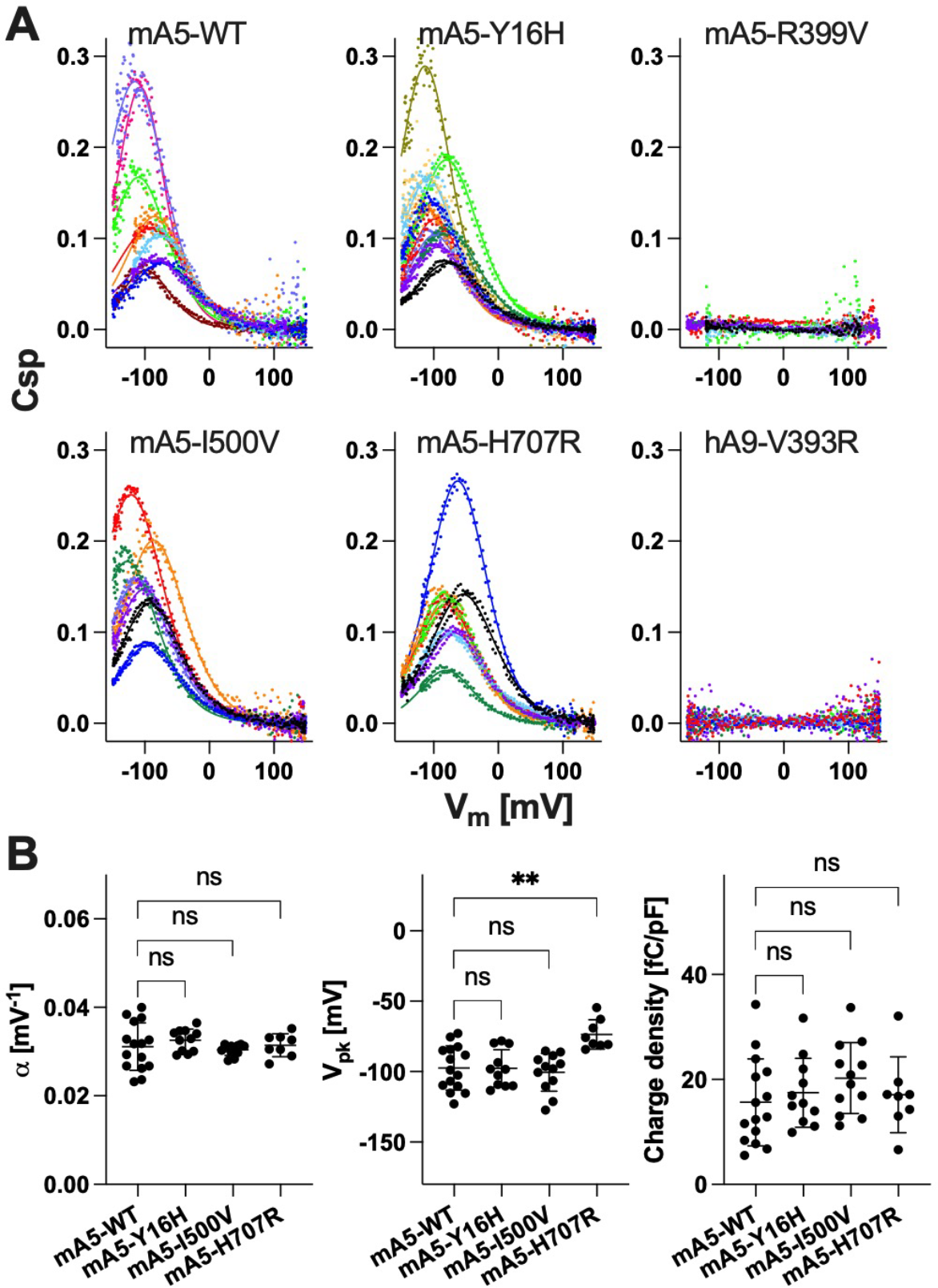
Whole-cell NLC recordings. (**A**) Representative NLC data. The magnitudes of NLC (C_m_ - C_lin_) were corrected for cell size (C_lin_), i.e., C_sp_ ≡(C_m_ - C_lin_)/C_lin_. Different colors indicate individual recordings. A two-state Boltzmann model was used to interpret the NLC data (solid lines). (**B**) Summaries of the NLC parameters. One-way ANOVA combined with the Tukey’s post hoc test was performed. “ns”, *p* ≥ 0.05. “**”, 0.001≤ *p* < 0.01.

### N- and C-terminal interactions in the cytosolic domains

H723R (c.2168A>G) is one of the most frequently found *SLC26A4* variants in patients, and preceding functional studies demonstrated that this common missense variant is dysfunctional (Ishihara et al., 2010; Jung et al., 2016; Lee et al., 2014; Wasano et al., 2020; Yoon et al., 2008; Zhang et al., 2022). Multiple other disease-associated missense variants affecting His^723^ in SLC26A4 have also been reported (Dai et al., 2008; Miyagawa et al., 2014; Van Hauwe et al., 1998), suggesting the importance of this His residue that is conserved among most SLC26 proteins (**Fig. 1B**). In the tertiary structure, this His residue is positioned very close to a Tyr residue at the N-terminus (Tyr^20^ in SLC26A4) (**Fig. 1**). Intriguingly, a disease-associated *SLC26A4* missense variant affecting this Tyr residue, Y20H (c.58T>C), was also reported (Zhao, 2014). The cryo-EM structures of SLC26A4, A5, A6, and A9 point to the possibility that these conserved His and Tyr residues may be involved in H-bond networks at the bottom of CD where the N-termini interact with C-termini (**Figs. 1C, 1D, and S1**). It is conceivable that these potential H-bond formations is dynamic and linked to the elevator-like large movements of the core domain to which the N-terminus is connected. Other residues that likely contribute to these H-bond networks are also conserved, and multiple *SLC26A4* missense variants affecting these residues have been reported to date, i.e., R24G (c.70C>G), R24L (c.71G>T), R24Q (c.71G>A), Y530H (c.1588T>C), Y530S (c.1589A>C), T721K (c.2162C>A), T721M (c.2162C>T), D724G (c.2171A>G), D724N (c.2170G>A), and D724V (c.2171A>T) (Anwar et al., 2009; Blons et al., 2004; Coyle et al., 1998; Prasad et al., 2004; Pryor et al., 2005; Rendtorff et al., 2013; Tian et al., 2021; Usami et al., 1999; Zhao et al., 2014). Among these variants, severe functional effects were demonstrated for Y530H, Y530S, T721M, and D724G (Choi et al., 2009; Ishihara et al., 2010; Pera et al., 2008; Wasano et al., 2020).

Using the fluorometric bicarbonate/chloride antiport assay that was established in our previous study (Wasano et al., 2020) with subsequent correction for protein expression in the cell membrane as was done for the iodide transport assay in **Fig. 4**, we found that H723D and H723Y severely impaired the function of SLC26A4 (**Fig. 6**), similarly to the common H723R. Y20H also resulted in the reduction of SLC26A4 antiport activity (**Fig. 6**). However, equivalent missense mutations only partially or barely affected the functions of SLC26A5 and SLC26A9. Specifically, Y16H did not alter the voltage sensitivity (α), voltage operating point (V_pk_), and charge density of SLC26A5 (**Fig. 5A**, top middle, and **5B**). H707R slightly shifted V_pk_ of SLC26A5, but α and charge density remained WT-like (**Fig. 5A** bottom middle, and **5B**). Y8H reduced the anion transport activity of SLC26A9 (**Figs. 3 and 4**), but its negative functional impacts were relatively milder compared to that of Y20H in SLC26A4 (**Fig. 6**). Lastly, the impact of H731R on the anion transport activity of SLC26A9 was very small (**Figs. 3 and 4**), which was in stark contrast to the severe function effect of H723R in SLC26A4 (**Fig. 6**).

**Figure 6.**
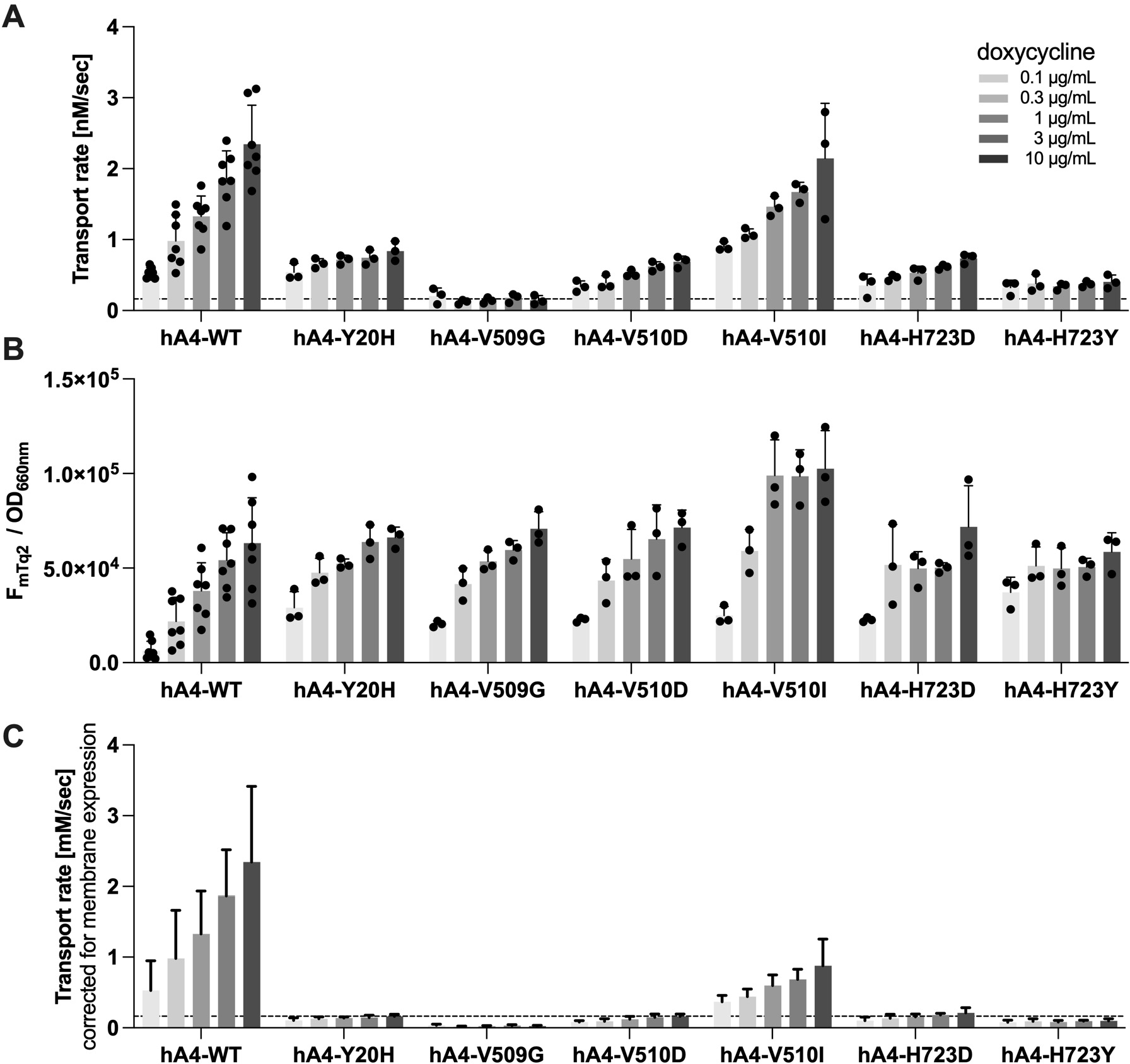
Multi-cell fluorometric bicarbonate/chloride antiport assays. (**A**) Bicarbonate/chloride antiport rates were measured in HEK293T cells expressing various SLC26A4 constructs in a Dox-dependent manner. A horizontal dashed line indicates the basal transport rate of non-induced cells (negative control). Error bars indicate standard deviation. (**B**) Ratios of mTq2 fluorescence (F_mTq2_) and the optical densities (OD_660nm_) of the cells expressing the mTq2-tagged SLC26A4 constructs, which quantify the total expressions of the SLC26A4 constructs. Error bars indicate standard deviation. (**C**) SLC26A4-mediate bicarbonate/chloride antiport rates were corrected for the relative membrane expressions as in Fig. 4.

### The homodimerization interface at the C-terminus of TM14

V499G is one of the artificial missense changes introduced to the junction between the TMD and the CD in SLC26A5 (Zheng et al., 2005). This mutation does not impair the membrane targeting of SLC26A5 but virtually eradicates the motor function of SLC26A5 within the physiologically relevant voltage range (Dallos et al., 2008; Homma et al., 2013). The cryo-EM structures of SLC26A5 revealed that Val^499^ and Ile^500^ are at the homodimerization interface at the C-termini of TM14 helices (Bavi et al., 2021; Butan et al., 2022; Futamata et al., 2022; Ge et al., 2021) (**Fig. 1**), implying the importance of these hydrophobic residues for stabilizing the homodimeric structure. To examine the general functional importance of the hydrophobic interaction between TM14 helices, we introduced V509G and V493G mutations to SLC26A4 and SLC26A9, respectively, which are equivalent to V499G in SLC26A5. We found that V509G completely abolishes the anion transport function of SLC26A4, whereas V493G only partially or barely affected chloride and iodide transport, respectively, in SLC26A9 (**Figs. 3 and 4**).

Like V499G, I500G was also shown to abolish the motor function of SLC26A5 (Futamata et al., 2022). Val^510^ in SLC26A4 is equivalent to Ile^500^ in SLC26A5, and a disease-associated missense change, V510D (c.1529T>A), has been reported (Jang et al., 2014). We found that this missense change also severely impairs the anion transport function of SLC26A4 (**Fig. 6**). The Val vs. Ile difference is trivial if any for SLC26A5’s motor function because I500V did not affect α, V_pk_, and charge density of SLC26A5 (**Fig. 5A**, bottom left, and **5B**). Interestingly, however, V500I in SLC26A4 largely reduced its anion transport function (**Fig. 6**).

Collectively, these observations imply the importance of the conserved hydrophobic residues at the C-termini of TM14 helices for fixing or confining the angles of the gate domains within a very small variable range in the cell membrane so that relative motions of the core domains with respect to the gate domains can complete alternating inward/outward-open conformational cycles in SLC26A4 or effectively translated into lateral expansion for electromotility in SLC26A5. It is conceivable that the uncoupled fast channel-like anion transport by SLC26A9 may not require fully completed elevator-like motion cycles.

### Attenuation of the inter-TM14 helices hydrophobic interaction induces hysteresis in SLC26A5 motor function

Mammalian SLC26A5 senses changes in transmembrane electric potentials (receptor potentials) and exerts electromotility, which is the molecular basis of cochlear amplification. It is likely that displacements of the core domains with respect to the gate domains (Bavi et al., 2021; Futamata et al., 2022; Ge et al., 2021; Kuwabara et al., 2023) are driven by voltage changes, and that NLC-inducing movements of a yet to be fully identified voltage sensing charge(s) or dipole(s) are coupled to the core domain displacements. As mentioned above, Val^499^ and Ile^500^ at the C-terminus of TM14 helix hydrophobically interact with Val^499^ and Ile^500^ in the other TM14 helix in the homodimeric structure. We found that slight attenuation of this hydrophobic interaction by I500V did not affect NLC (see above). However, further attenuation of the hydrophobicity by I500A induces hysteresis in NLC measured by a sinusoidal voltage protocol (**Fig. 7A**) in mouse (**Fig. 7B**), gerbil (**Fig. 7C**), and naked mole-rat SLC26A5 (**Fig. 7D**). The observed NLC hysteresis can be explained by hyperpolarization- and depolarization-induced changes in the angle between the two TM14 helices, which would change the direction of the core domain displacements and concomitantly change the vector of the voltage-sensing charge movement. Thus, the emergence of NLC hysteresis supports the idea that SLC26A5’s voltage sensor charge movement is tightly coupled to the relative displacements of the core domains with respect to the gate domains (**Fig. 7E**).

**Figure 7.**
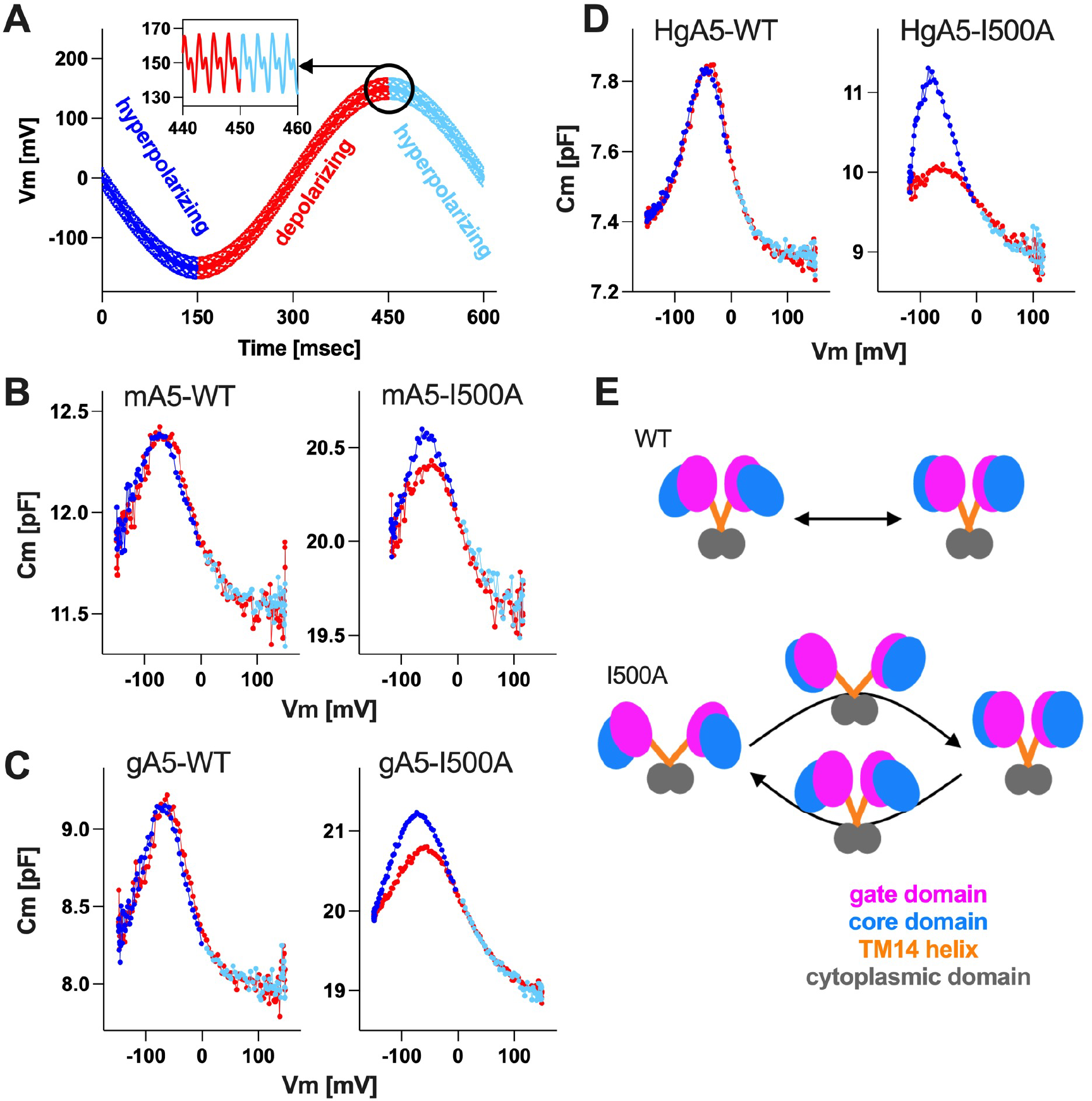
NLC hysteresis induced by I500A in SLC26A5. (**A**) The voltage protocol used for whole-cell NLC recordings. An inset shows dual sinusoidal stimuli (390.6 and 781.2 Hz, 10 mV) superimposed onto a large sinusoidal voltage stimulus (with either 120 or 150 mV amplitude). (**B-D**) NLC recorded in HEK293T cells expressing WT or I500A of mouse (mA5, panel B), gerbil (gA5, panel C), and naked mole-rat (HgA5, panel D) SLC26A5. C_m_ values determined by hyperpolarizing and depolarizing voltage stimulus phases are indicated in different colors and matched with the color coding used in panel A. (**E**) A hypothetical model to account for the observed NLC hysteresis (see the main text).

## DISCUSSION

The homodimeric structures of SLC26A4, A5, A6, and A9 are strikingly similar. Given the high degree of the amino acid sequence similarity, it is probable that the other SLC26 family proteins also have similar homodimeric structures. Elevator-like motions of the core domains with respect to the gate domains intuitively account for the anion transport mechanism. In fact, inward- and outward-open conformations were experimentally captured for SLC26A4 (Liu et al., 2023). This gate/core relative motions can also account for the voltage-driven motor function of mammalian SLC26A5 (Bavi et al., 2021; Ge et al., 2021; Kuwabara et al., 2023), implying that voltage-sensing mechanism in SLC26A5 may not be unique in the SLC26 family. In fact, SLC26A4 also exhibits a large NLC but with a shallow voltage sensitivity (small α) and extremely hyperpolarized V_pk_ (Kuwabara et al., 2018). It is likely that movements of a voltage sensing charge(s) and/or dipole(s) are tightly coupled to the movements of the core domains, and that the efficiency of this electromechanical coupling is evolutionary enhanced and optimized in mammalian SLC26A5. It is curious to learn if the loss of anion transport function is inevitable for optimizing the electromechanical coupling efficiency in mammalian SLC26A5. Comparative structural and functional studies on nonmammalian transporter SLC26A5 may address this outstanding question.

Whether the protomers functionally communicate in a homodimer complex is another outstanding question. The N- and C-termini are connected to the core and the gate domains, respectively, and interwound with one another at the bottom of the CD (**Figs. 1 and S1**). Previous studies demonstrated the importance of these N- and C-terminal regions for the function of SLC26A5 (Navaratnam et al., 2005; Zheng et al., 2005). In SLC26A5, we presume that the N- and C-terminal interaction is important for stabilizing homodimerization, but not for inter-protomer communication, because coexpression of a dysfunctional SLC26A5 (V499G/Y501H) did not affect the function of WT SLC26A5 upon heteromerization (Homma et al., 2013). Consistently, the present study found that Y16H and H707R barely affect NLC of SLC26A5 (**Fig. 5**). On the contrary, missense changes at equivalent sites in SLC26A9 (Y8H and H731R) and SLC26A4 (Y20H, H723D, H723Y) partially reduced or completely abrogated the anion transport functions, respectively (**Figs. 3, 4, and 6**). Based on these observations, we speculate that inter-protomer functional interaction indeed exists in transporter SLC26 proteins and is particularly crucial for coupled anion transport such as bicarbonate/chloride exchange mediated by SLC26A4. The presence of many disease-associated *SLC26A4* missense variants that potentially affect the N- and C-terminal interaction is in line with such speculation.

We showed that V393R abolished the anion transport function of SLC26A9 (**Figs. 3 and 4**), but this single missense change did not confer NLC on SLC26A9 (**Fig. 5**). Likewise, R399V abolished NLC of SLC26A5 (**Fig. 5**), but it conferred only a very small chloride conductance on SLC26A5 (**Fig. 3**). It thus appears that the diverse functions of the SLC26 proteins are not attributable to a single or a few local amino acids change. Despite the overall structural similarity, the packing of the fourteen TM helices differs among the SLC26 family of proteins (**Fig. 1**). The slightly denser packing of TM helices in mammalian SLC26A5 may be crucial for maximizing the electromechanical coupling efficiency and reducing the energy barrier between an expanded and a compacted states for attaining fast conformational transition kinetics (i.e., fast electromotility). This speculative optimization may have resulted in loss of anion transport function during molecular evolution but may have been tolerated because the anion transport activity of SLC26A5 is of minimal physiological significance in mammalian hearing (Takahashi et al., 2023). Again, comparative structural and functional studies on nonmammalian transporter SLC26A5 may provide definitive mechanistic insights as to how the fast and efficient electromechanical coupling in mammalian SLC26A5 has evolved from ancestral anion transporters.

Although this study focused only on the SLC26 molecule themselves for molecular mechanisms, other factors, e.g., interacting proteins, also play roles to support their diverse physiological functions. For example, apical vs. basolateral membrane localization of SLC26 proteins, which is probably dictated by yet to be fully defined distinct protein interactomes (Cimerman et al., 2013; Li et al., 2020; Sengupta et al., 2010), is of obvious physiological significance but was not dealt in the present study using the nonpolarized cell line, HEK293T. Regulatory interactions are also possible (Bai et al., 2010; Chang et al., 2009a; Homma et al., 2010; Keller et al., 2014; Ko et al., 2004; Ohana et al., 2013; Xu et al., 2022) but not controlled, either, in the present study. It is likely that intrinsically disordered regions that were not resolved in the SLC26 structures reported to date are responsible for determining distinct membrane targeting and regulatory interactions, which need to be fully addressed in future studies. Nevertheless, the knowledge obtained by this comparative functional study would be useful in predicting the pathogenicity of variants found in this functionally diverse family of proteins.

## METHODS

### Generation of stable cell lines that express various SLC26 protein constructs

cDNAs encoding human SLC26A4, mouse SLC26A5, gerbil SLC26A5, naked mole-rat SLC26A5, human SLC26A9 (WT and missense mutants) with a C-terminally attached mTurquoise (mTq2) tag were cloned into a pSBtet-pur vector (Addgene) using SfiI sites. Stable cell lines expressing these SLC26 constructs in a doxycycline-dependent manner were established in HEK293T cells as previously described (Wasano et al., 2020). Stable cells were maintained in DMEM supplemented with 10% FBS and 1μg/ml puromycin (Fisher Scientific).

### Whole-cell recordings

Whole-cell recordings were performed at room temperature using the Axopatch 200B amplifier (Molecular Devices) with a 10 kHz low-pass filter. Recording pipettes pulled from borosilicate glass were filled with an ionic blocking intracellular solution containing (mM): 140 CsCl, 2 MgCl_2_, 10 EGTA, and 10 HEPES (pH 7.4). L-aspartate was used to replace chloride for preparing low chloride-containing intracellular solutions. Cells were bathed in an extracellular solution containing (mM): 120 NaCl, 20 TEA-Cl, 2 CoCl_2_, 2 MgCl_2_, 10 HEPES (pH 7.4). Osmolality was adjusted to 309 mOsmol/kg with glucose. Command voltages were step functions of 30 msec duration (from -100 mV to +100 mV or -50 mV to +50 mV, in 10 mV steps). Holding potentials were set to 0 mV. NLC was measured using sinusoidal voltage stimuli (2.5-Hz, 120 or 150 mV amplitude) superimposed with two higher frequency stimuli (390.6 and 781.2 Hz, 10 mV amplitude). Data were collected by jClamp (SciSoft Company, New Haven, CT) (Santos-Sacchi et al., 1998).

### NLC data analysis

Voltage-dependent C_m_ data were analyzed using the following two-state Boltzmann equation:

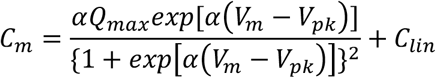

where α is the slope factor of the voltage-dependence of charge transfer, Q_max_ is the maximum charge transfer, V_m_ is the membrane potential, V_pk_ is the voltage at which the maximum charge movement is attained, and C_lin_ is the linear capacitance. The specific capacitance, C_sp_, was calculated as (C_m_ – C_lin_)/C_lin_.

### Fluorometric anion transport assay

Fluorometric HCO_3-_/Cl^-^ and I^-^/Cl^-^ antiport assays were established and described in detail in a previous study (Wasano et al., 2020). Briefly, stable HEK293T cells expressing SLC26 protein constructs or those with coexpressed iodide sensitive fluorescent protein, mVenus^H148Q/I152L^, were used for HCO_3-_/Cl^-^ and I^-^/Cl^-^ antiport assays, respectively. For the HCO_3-_/Cl^-^ antiport assay, cells were loaded with a pH indicator, SNARF-5F (S23923, Thermo Fisher Scientific) in a high chloride buffer containing (mM): 140 NaCl, 4.5 KCl, 1 MgCl_2_, 2.5 CaCl_2_, 20 HEPES (pH 7.4). The antiport assay was initiated by an automated injection of a low chloride buffer containing (mM): 125 Na-gluconate, 5 K-gluconate, 1 MgCl_2_, 1 CaCl_2_, 20 HEPES, 25 NaHCO_3_ with 5% CO_2_ in Synergy Neo2 (Agilent/BioTek). For the I^-^/Cl^-^ antiport assay, cells were resuspended in a high chloride buffer containing (mM): 150 NaCl, 1 MgCl_2_, 1 CaCl_2_, 20 HEPES (pH 7.5). The I^-^/Cl^-^ antiport assay was initiated by an automated injection of a high iodide buffer containing (mM): 150 NaI, 1 MgCl_2_, 1 CaCl_2_, 20 HEPES (pH 7.5). The fluorescence of SNARF-5F (for HCO_3-_/Cl^-^ antiport assay) or mVenus^H148Q/I152L^ and mTq2 (for I^-^/Cl^-^ antiport assay) were measured in a time dependent manner using Synergy Neo2 (Agilent/BioTek) and the data analyzed offline as described previously (Wasano et al., 2020).

### Cell surface protein labeling and quantitation

Stable cells expressing SLC26 constructs were washed once with PBS, and 10 μM Sulfo-Cyanine3 NHS ester (Lumiprobe) dissolved in 2 mL of ice-cold PBS (per well of 6-well plate) was added and incubated for 30 minutes at 4 °C. The reaction was stopped by the addition of 200 μL of 100 mM glycine (per well). Cells were collected and lysed by sonication on ice in 500 μL of a lysis buffer containing 150 mM NaCl, 20 mM HEPES, pH 7.5, 1 mM EDTA, 20 mM DDM, 1 mM DTT, and 50 μg/mL leupeptin. The lysate was centrifuged at 16,000 × *g* for 5 minutes at 4 °C. A GFP selector slurry (5 μL, NanoTag Biotechnologies) was added to the supernatant and incubated for 30 minutes at 4 °C, with end-over-end mixing using a rotator. Bound proteins were collected alongside the GFP selector by brief centrifugation, and observed with a fluorescent microscope (Leica DMIRB). Merged images of GFP selectors in cyan and red channels were analyzed using Fiji (Schindelin et al., 2012) to determine the fluorescent signal intensities of Cyanine3 (F_Cy3_) and mTq2 (F_mTq2_).

### Propagated error calculation

Error propagations (σ) were calculated by the following equations:

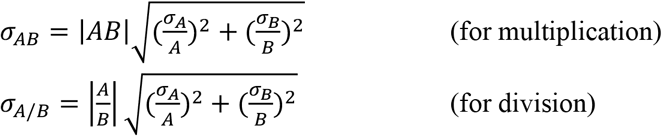

where A and B are the mean values with associated errors, α _A_ and α _B_, respectively.

### Statistical analyses

Statistical analyses were performed using Prism (GraphPad software). One-way analysis of variance (ANOVA) combined with Tukey’s posttest was used for multiple comparisons. *p* < 0.05 was considered statistically significant.

## ACKNOWLEDGEMENTS

This work was supported by an NIH grant DC017482 (to KH) and by the Hugh Knowles Center.

## AUTHOR CONTRIBUTIONS

K.H. initiated the study. S.T. and K.H. performed experiments, analyzed the data, and wrote the manuscript.

## FIGURES AND LEGENDS

**Figure S1.**
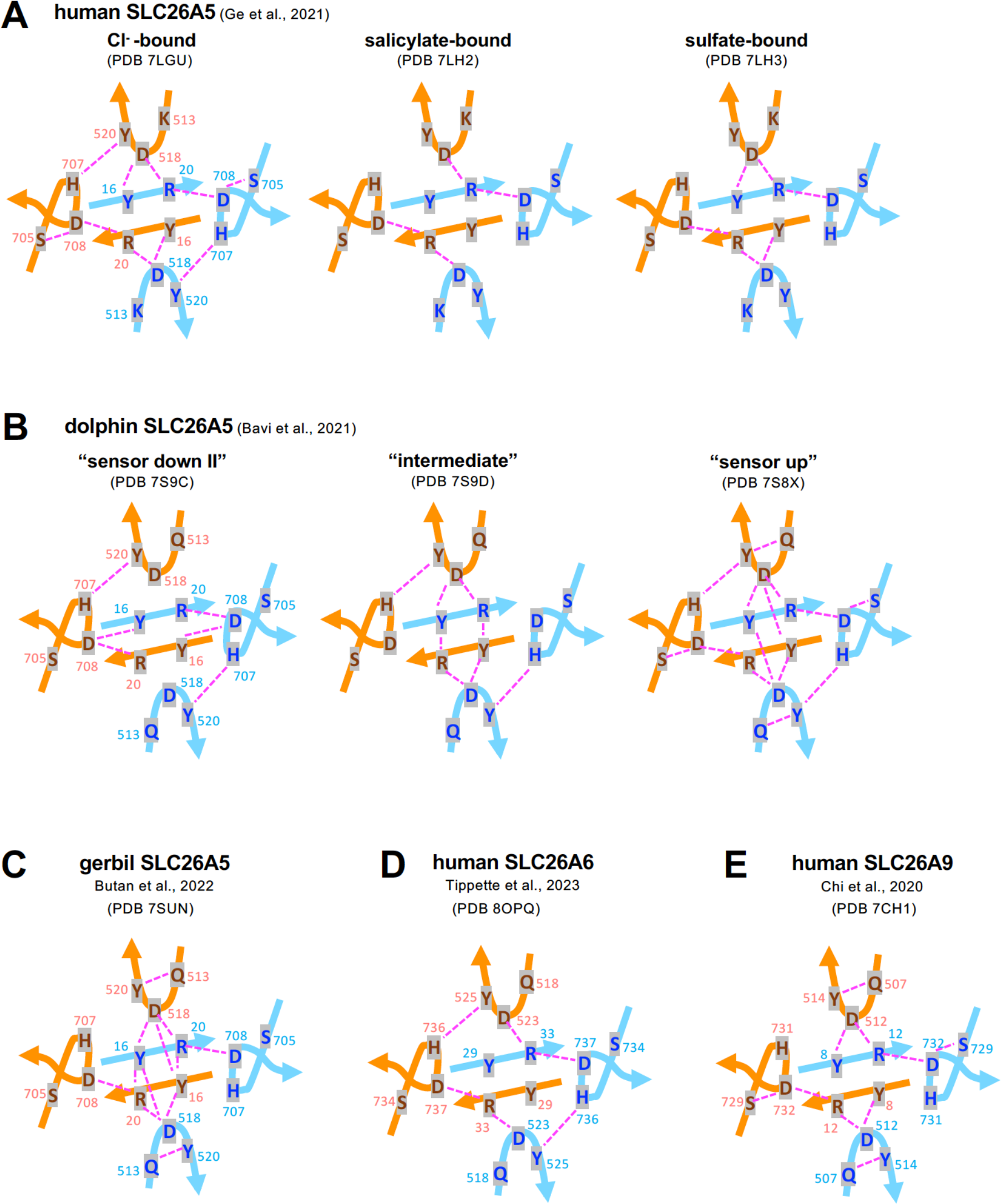
Schematic representations of the N- and C-terminal interactions in human SLC26A5 (A), dolphin SLC26A5 (B), gerbil SLC26A5 (C), human SLC26A6 (D), human SLC26A9 (E). Magenta dashed lines indicate potential hydrogen bond interactions. This supplementary figure is related to Fig. 1 in the main text.

